# Probing the sequence variability tolerance in a de novo α-helical barrel biocatalyst

**DOI:** 10.64898/2026.09.28.754888

**Authors:** Wael Elaily, Markus Braun, David Stoll, Morakot Chakatok, Matteo Aleotti, Birgit Grill, Aleksandar Bijelic, Mélanie Hall, Gustav Oberdorfer

## Abstract

De novo-designed enzymes have recently achieved high catalytic activity and stereoselectivity while demonstrating exceptional thermostability in entirely novel protein scaffolds. Among these, α-helical barrel protein scaffolds are attractive structures for biocatalysis due to their structural simplicity, high thermostability, and rationalizable sequence patterning. However, enabling major structural reengineering of these scaffolds while maintaining the structure, stability and catalytic activity while also improving soluble protein production remain major challenges and pose the fundamental question how engineerable a de novo backbone-sequence pair is. Here, we combine deep learning based and classic computational protein design to modify and optimize de novo α-helical barrel biocatalysts. Using the previously reported six-helical barrel 6H5L as a model scaffold, AlphaFold2-guided RosettaRemodel enabled the design of a truncated variant, whose crystal structure closely matches the design model. Additional sequence-redesign using ProteinMPNN generated a variant with a tenfold increase of soluble protein yield in Escherichia coli. Biochemical, biophysical, and structural analyses showed that both variants retained the overall barrel architecture, high thermal stability, and catalytic activity for both purified protein and whole-cell systems. Detailed kinetic analysis on the variants showed both variation in k_cat_ and K_m_, reflecting changes in catalytic turnover and substrate binding. Together, these approaches provide new insights and possibilities for the further engineering of functional de novo α-helical barrels, their ability to withstand dramatically large sequence changes and their broader application in biocatalysis and biotechnology.

## Introduction

Enzymes are remarkable catalysts due to their exceptional efficiency, selectivity, and activity under benign reaction conditions, making them attractive alternatives to traditional chemical catalysts. However, natural enzymes, evolved for metabolic reactions, face several challenges for broader industrial use, including low production yields, limited stability across hosts, and poor activity under industrially relevant reaction conditions^1^.

Computational enzyme design offers a promising route to overcome these limitations through innovative protein design strategies^2–4^. Both reengineering existing enzymes and creating new de novo catalysts have yielded proteins capable of catalyzing non-natural reactions with high efficiency and selectivity, demonstrating the transformative potential of computational protein design in biocatalysis ^5–7^.

Beyond traditional physics-based methods such as Rosetta^8,9^, advances in deep learning (DL)-based techniques^10–14^, including Protein Message Passing Neural Network (ProteinMPNN)^15^ for sequence design, have expanded the scope of protein engineering. DL tools complement structure prediction models, such as AlphaFold2^16^ (AF2), RoseTTAFold2^17^, ESMFold^18^, and OmegaFold^19^, effectively linking sequence generation to structure prediction. Integrating these methods increases design accuracy and efficiency, enabling flexible workflows that combine DL and physics-based techniques. Among these, ProteinMPNN has demonstrated strong computational and experimental performance, improving soluble expression, stability, and activity of redesigned proteins ^15^ while maintaining structural consistency with predicted model. This synergy suggests broad potential for ProteinMPNN across diverse enzyme families and justifies experimental exploration.

A representative application involves de novo (retro)-aldolases, whose well-understood lysine-dependent mechanism, characteristic of class I aldolases^20,21^, makes them valuable model systems. Artificial aldolases have been developed using diverse scaffolds, including TIM-barrels^2,5,22,23^, β-barrels^24^, mini-proteins^25^, and, more recently, alpha-helical scaffolds^26^ and helical barrels^27^. This structural diversity establishes aldolase reactions as ideal benchmarks for evaluating DL–based protein design strategies.

Building on this foundation, catalytically active de novo α-helical bundle proteins are emerging as promising scaffolds due to their inherent thermodynamic stability^27,28^, solvent resistance, and structural simplicity^27,29^. These characteristics make them particularly suitable for the development of robust and adaptable biocatalysts capable of functioning under demanding conditions^27^.

In this study, we focused on the question how much sequence variation and structural diversification the catalytically active de novo six-helical barrel 6H5L can withstand and if it is possible to optimize its recombinant expression. The structure was originally designed parametrically with Rosetta and optimized for (retro)-aldol catalysis^27^. Two complementary strategies were pursued. First, we used ProteinMPNN together with AF2 to redesign the 6H5L_RA1 variant—previously reported to exhibit a 3.5-fold higher catalytic activity than the original 6H5L^27^, to enhance soluble expression. Second, we employed RosettaRemodel^30^ guided by AF2 to generate a truncated 6H5L variant that retained its thermostability, catalytic activity, and barrel architecture.

Together, these approaches yielded two new de novo protein designs, and their experimental characterization allowed us to evaluate the predictive accuracy of AF2 for outputs generated using both physics-based (RosettaRemodel) and DL-driven (ProteinMPNN) methods. By focusing on α-helical de novo scaffolds as both input and output, this work provides insights into the combined potential of DL and physics-based design for developing robust, high yield, and structurally diverse de novo biocatalysts for applications in sustainable and green chemistry.

## Results

### Design strategies

The strategy based on ProteinMPNN to redesign the 6H5L_RA1 model, an optimized variant of 6H5L with a 3.5-fold higher retro-aldolase activity^27^, involved a complete redesign of the bundle surface and the unoptimized cavity **1** in the hydrophobic channel, while keeping the previously optimized cavity **2** fixed (Figure 1A). From the resulting outputs, we selected two models based on high AF2 pLDDT scores and distinct isoelectric points: 6H5L_mpnn1740 (pI 5.4) and 6H5L_mpnn1409 (pI 7.18). Both represent alternative long-barrel variants selected for experimental characterization of their expression, structural properties, and catalytic performance.

**Figure 1.**
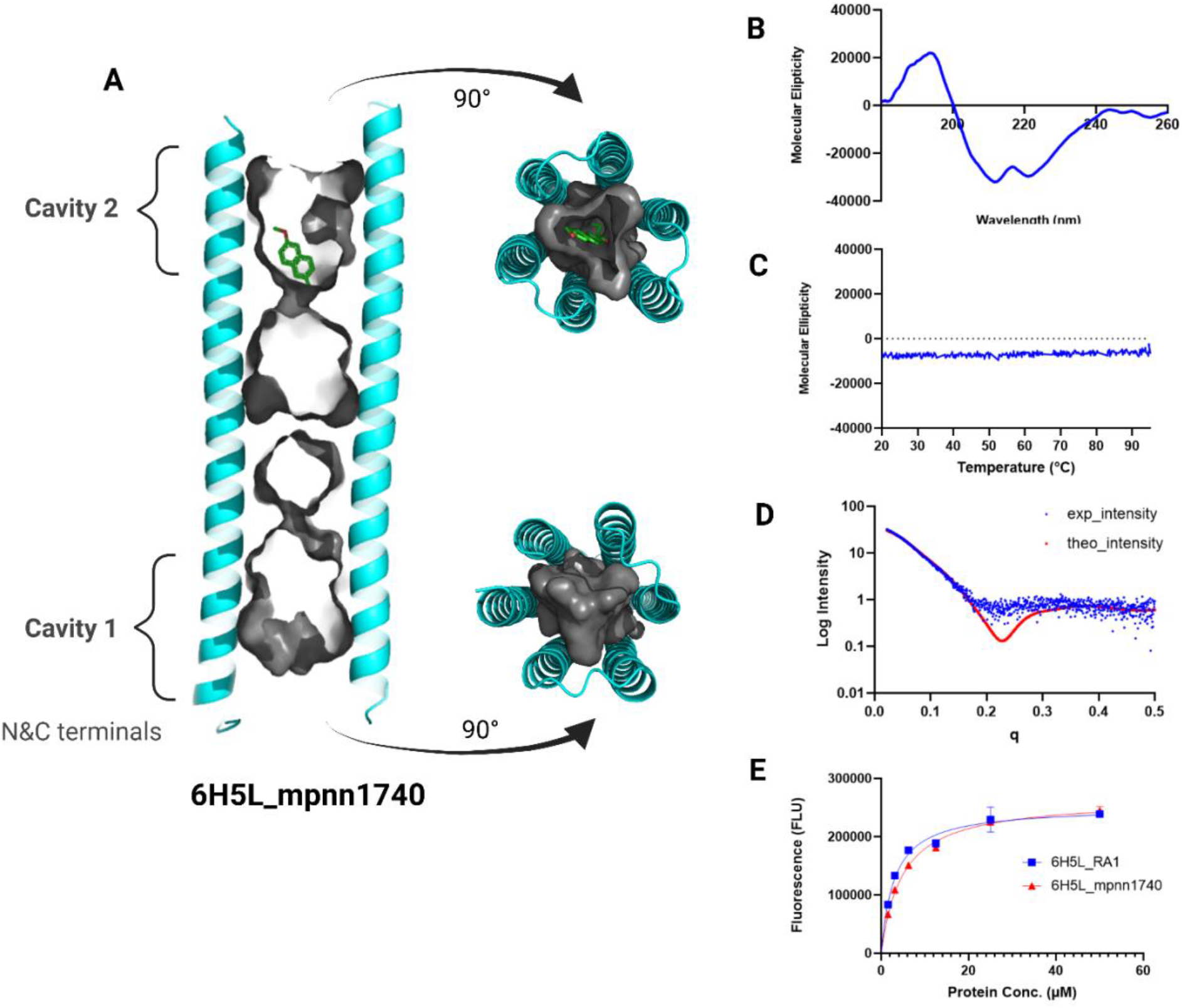
Structural characterization of 6H5L_mpnn1740 design. **A)** AlphaFold2 (AF2) predicted structure (cyan) shows two cavities at both termini of the helix bundle: cavity 1 (unoptimized) and cavity 2 (optimized and including the substrate in its diketone form **5**). **B)** Circular dichroism (**CD**) spectrum confirming a predominantly α-helical structure. **C)** Temperature scan profile demonstrating high thermal stability with only minor changes up to 95 °C. **D) SAXS** scattering data and fit between the theoretical (red) and measured (blue dots) data showing an excellent agreement; chi^2^ = 2.5. **E) DPH assay** showing saturation binding curves of 6H5L_mpnn1740 (red triangles) vs. input design 6H5L_RA1 (blue squares) at different protein concentrations and constant DPH concentration (1 μM).

To complement these redesign efforts and further examine how barrel length and electrostatic distribution influence structural stability and catalytic potential, we employed RosettaRemodel^30^ to create a truncated version of the 6H5L scaffold. The resulting designs yielded shorter barrels open at both termini, retaining only one of the two charged rings in the hydrophobic channel of 6H5L (each containing three lysine and three complementary glutamate residues, Figure 2A)^27^. By altering the cutting site, we obtained two variants: 6H5L_Cut with 180 amino acids and 6H5L_Cut2 with 162 amino acids. Both showed high AF2 pLDDT scores (>90 and >80, respectively), although 6H5L_Cut2 exhibited a lower mean confidence (~60) across all five AF2 models (Figure S2). These constructs enabled testing of AF2 accuracy in predicting de novo α-helical bundles generated by RosettaRemodel^30^. This approach also enabled the development of a compact six-helix barrel with two open cavities connected through the hydrophilic residues of ring 1 (Figure 2A), offering potential for more complex or coenzyme-dependent catalytic reactions.

**Figure 2.**
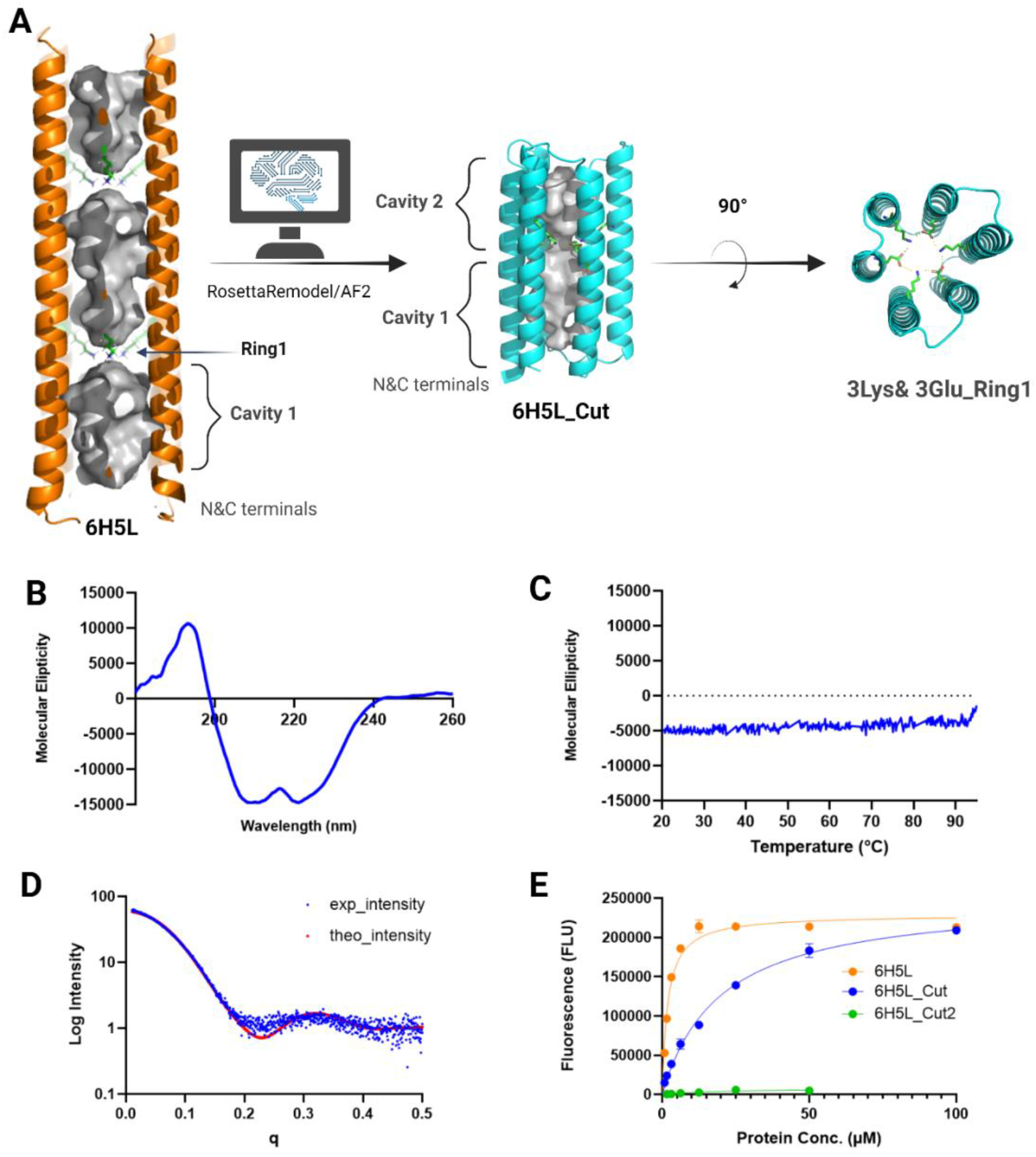
Design scheme and structural characterization of the 6H5L truncated designs. **A)** Scheme illustrating the design approach of the truncated versions. The AlphaFold2 (AF2) structure of 6H5L (orange) vs. that of the truncated version 6H5L_Cut (cyan), with two cavities formed by introducing ring **1** in the hydrophobic channel (each containing three lysine residues paired with three glutamate residues). **B)** Circular dichroism (**CD**) spectrum of 6H5L_Cut design confirming a predominantly α-helical structure. **C)** Temperature scan profile of 6H5L_Cut design demonstrating high thermal stability with only minor changes up to 95 °C. **D) SAXS** scattering data of 6H5L_Cut and fit between the theoretical (red) and measured (blue dots) data showing an excellent agreement; chi^2^ = 2.4. **E) DPH assay** showing saturation binding curves of 6H5L_Cut vs. input design 6H5L at different protein concentrations and constant DPH concentration (1 μM), as well as the misfolded design (6H5L_Cut2).

### Soluble protein production

The His-tagged designs were produced in *E. coli* and purified using Ni-NTA affinity chromatography, with each step monitored by SDS-PAGE. Among the MPNN variants, 6H5L_mpnn1740 (pI 5.4) showed strong soluble expression at ~35 kDa (with theoretical MW = 34.82 kDa), comparable to the reference 6H5L_RA1 design (Figure S1), whereas 6H5L_mpnn1409 (pI 7.18) displayed a faint band, indicating weaker expression. Size-exclusion chromatography (SEC) of 6H5L_mpnn1740 revealed a single, intense peak at the same elution volume as 6H5L_RA1, indicating a comparable monomeric state. Notably, 6H5L_mpnn1740 showed a tenfold higher protein yield, reaching 16.7 mg/L compared with 1.7 mg/L for 6H5L_RA1. In contrast, 6H5L_mpnn1409 produced multiple peaks, suggesting reduced stability or partial misfolding (Figure S1).

For the truncated variants, 6H5L_Cut and 6H5L_Cut2, SDS-PAGE showed strong bands at ~20 kDa (with theoretical MW = 18.19 kDa). The SEC profile of 6H5L_Cut (with theoretical MW = 21.03 kDa) showed a monodisperse peak, whereas 6H5L_Cut2 exhibited multiple peaks, consistent with aggregation and impurities (Figure S2). To ensure the accuracy of these findings, all SEC fractions corresponding to major peaks were verified by SDS-PAGE and collected for further characterization.

### Structural characterization

#### DPH assay

To assess proper barrel folding, 1,6-diphenyl-1,3,5-hexatriene (DPH) fluorescence assays were performed using 1 µM DPH dye and protein concentrations up to 100 µM. As DPH fluorescence intensifies in hydrophobic environments, the binding curve reflects cavity integrity^31^. The MPNN variant 6H5L_mpnn1740 exhibited a saturation profile similar to 6H5L_RA1, but with approximately twice the *K*_D_ (Figure 1E, Table 1), likely due to partial obstruction of cavity 1 during redesign (Figure 1A). The truncated 6H5L_Cut variant showed distinctly altered binding-curve behavior, along with elevated K_D_ values, yet still within the micromolar range. (Figure 2E, Table 1), indicating weaker binding affinity. In contrast, 6H5L_Cut2, which showed multiple SEC peaks and low AF2 pLDDT scores (Figure S2), exhibited no detectable DPH binding (Figure 2E), suggesting misfolding and failure to form the expected six-helix barrel.

**Table 1.**
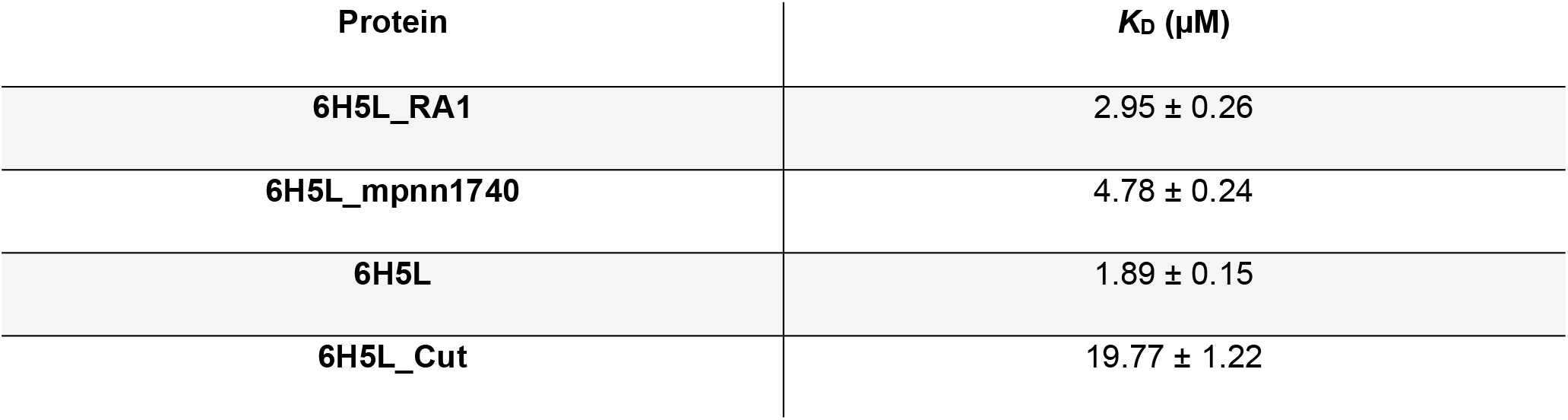
Dissociation constants (K_D_) for ligand-binding reactions of various protein variants.

#### Circular dichroism spectroscopy (CD) and temperature scan

Both 6H5L_mpnn1740 and the truncated version 6H5L_Cut exhibited CD spectra characteristic of α-helical proteins, with minima at 222 nm and 208 nm (Figures 1B and 2B). Temperature scans showed that both designs remained stable up to 95 °C (Figures 1C and 2C), consistent with previous reports for the parental 6H5L and the optimized variant 6H5L_RA1^27^.

#### Small-angle X-ray scattering (SAXS)

SAXS profiles of the 6H5L_mpnn1740 and 6H5L_Cut designs were obtained at protein concentrations of 3 and 6 mg/mL, respectively. The results for both proteins showed a good fit between the measured and calculated scattering profiles, indicated by chi^2^ values of 2.5 and 2.4, respectively (Figures 1D and 2D). The data confirm that both designs exhibit the expected α-helical barrel conformation in solution, consistent with AF2 predictions. Model fitting was performed using the FoXS online server ^32^.

#### X-ray Crystallography

The crystal structure of the truncated variant 6H5L_Cut (PDB: 9R2B) closely matches the AlphaFold-predicted design, with a C_α_ RMSD of 2.61 Å over 171 aligned residues (Fig. 3A). While the overall six-helix barrel architecture and open hydrophobic channel are fully preserved, the crystal structure reveals a noticeable deviation from the nearly symmetric, cylindrical geometry of the design (Fig. 3B). In particular, the individual helices exhibit differential changes in their orientation relative to the central bundle axis, resulting in an anisotropic deformation of the bundle. This structural shift is driven primarily by helix H5, which tilts by 6.3° relative to the bundle axis (vs. 1.9° in the model). As a result, the bundle expands along the H1–H4 axis (18.3 Å to 22.0 Å) and constricts along H2–H5 (21.6 Å to 17.7 Å), substantially increasing the anisotropy of the bundle geometry in the crystal structure compared with that of the predicted model. Despite this global geometric shift, the six charged residues lining the central cavity remain in nearly the same positions as designed, with a heavy-atom side-chain RMSD of 1.2 Å (Fig. 3C). This shows that the designed electrostatic interaction network is maintained in the crystal structure, preserving the barrel architecture while keeping the central cavity accessible for substrate binding and catalysis.

**Figure 3.**
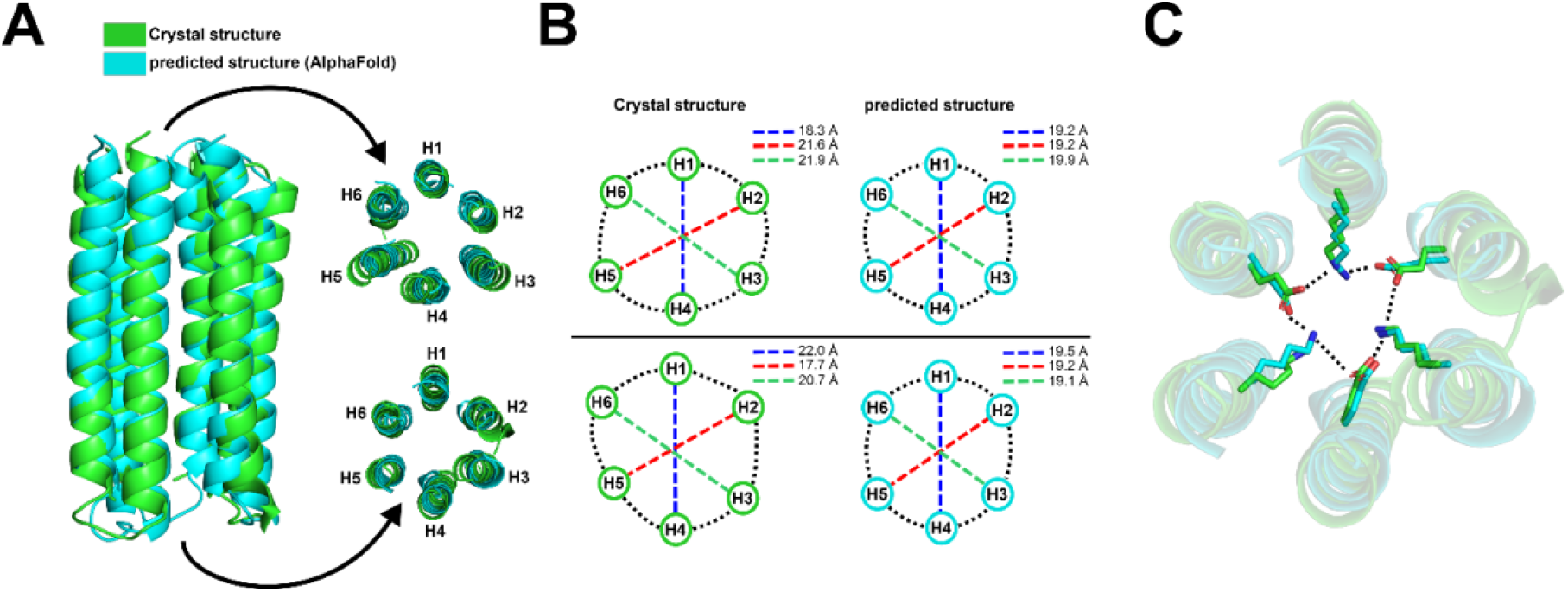
Comparison of the crystal structure of 6H5L_Cut (PDB: 9R2B) vs. the AlphaFold-predicted model. (A) Superimposition of the predicted model (cyan) and crystal structure (green). The end-on views on the right side show the relative arrangement of the α-helices H1–H6 at the two opposing ends of the bundle. (B) Geometric comparison of the bundle cross-sections at the top and bottom ends. Distances between opposing helix axes are shown (colored dotted lines). Black curved dotted lines indicate the shape of the overall cross-section of the bundle. (C) Close-up of the central cavity region, illustrating that the six charged residues are in nearly identical conformations in the crystal structure and predicted model, with side-chain RMSD of 1.2 Å.

### Biocatalytic activity and whole-cell biotransformation

#### Retro-aldol inhibition reaction

To confirm the presence of a catalytically active lysine residue in 6H5L_mpnn1740 and 6H5L_Cut, we performed an inhibition assay using a 1,3-diketone analog (5), which forms a covalent enamine with the catalytic lysine residue (H2N-Enz, Figure 4B). Both designs exhibited a new UV-Vis absorption peak at 340 nm compared to the free protein, confirming the formation of a vinylogous amide via a Schiff base (Figure 4A). These results confirm the preservation and integrity of the nucleophilic lysine residue and its ability to form a Schiff base with the diketone analog, a key step in the catalytic mechanism.

**Figure 4.**
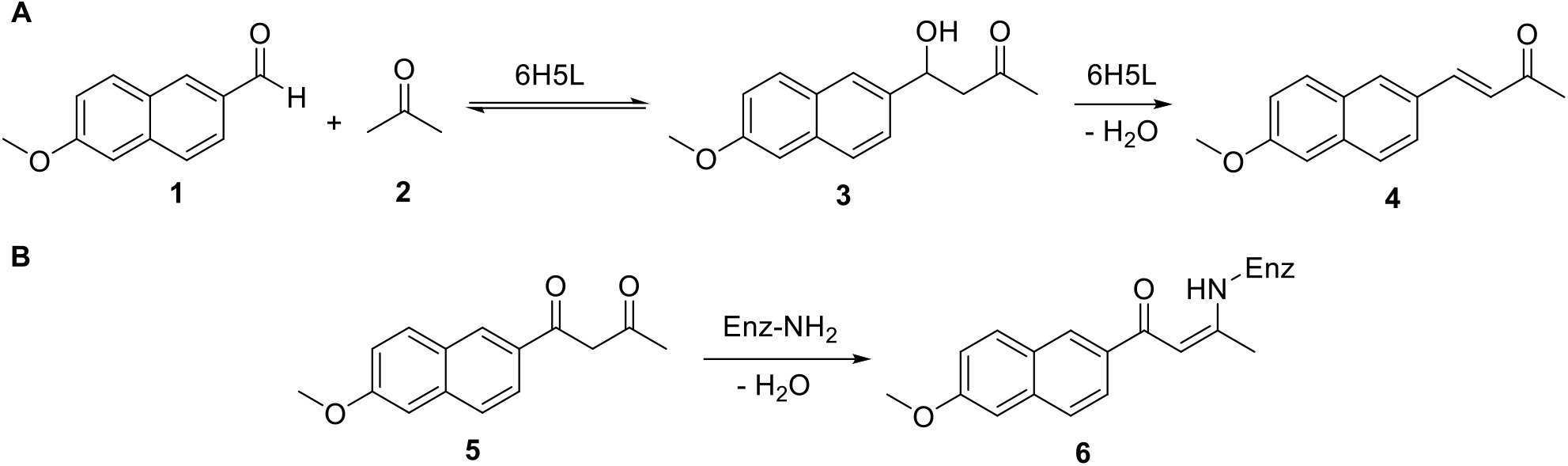
**A)** Scheme of the reversible aldol reaction of 6-methoxy-2-naphthaldehyde **1** and acetone **2** producing 4-hydroxy-4-(6-methoxy-2-naphthyl)-2-butanone (methodol **3**) and the corresponding *α*,β-unsaturated ketone **4. B)** Inhibition reaction by 1,3-diketone **5**, resulting in the formation of a vinylogous amide **6** covalently bound to the catalytic lysine residue (H_2_N-Enzy).

#### Activity assay

Retro-aldolase activity was measured using a fluorescence assay detecting the formation of 6-methoxy-2-naphthaldehyde 1 from racemic methodol 3 (Figure 5A)^33^. The 6H5L_mpnn1740 variant retained catalytic activity but showed a lower second-order rate constant (*k*_cat_/ *K*_m_ = 2.88 M^−1^ min^−1^) than the input design 6H5L_RA1 (5.09 M^−1^ min^−1^) (Figure 5B, Table 2). However, when looking at the individual kinetic parameters, 6H5L_mpnn1740 showed a higher *k*_cat_ than 6H5L_RA1 (48.45 × 10^−4^ vs. 27.31 × 10^−4^ min^−1^), while its *K*_m_ was approximately threefold higher (1580 vs. 530 µM). This suggests that the redesigned variant has a higher turnover rate but lower substrate affinity, resulting in lower overall catalytic efficiency. The increased *K*_m_ may be explained by partial obstruction of the unoptimized cavity 1 during ProteinMPNN redesign (Figure 1A).Comparison with the previously optimized 6H5L_RA2 variant, which possesses a single active pocket with a lysine at position K223, revealed similar kinetic parameters (Table 2), supporting this interpretation.

**Table 2.** Retro-aldolase kinetic parameters of all protein variants.

| PROTEINS | $K_M$<br>($\mu\text{M}$) | $K_{\text{cat}}$<br>($\times 10^{-4} \text{ min}^{-1}$) | $K_{\text{cat}}/K_M$<br>($\text{M}^{-1} \text{ min}^{-1}$) |
| --- | --- | --- | --- |
| 6H5L | 371±61 | 7.29±0.07 | 1.96 |
| 6H5L_Cut | 382±82 | 9.09±0.82 | 2.38 |
| 6H5L_RA1 | 530±50 | 27.31±1.19 | 5.09 |
| 6H5L_mpnn1740 | 1580±384 | 48.45±7.68 | 2.88 |
| 6H5L_RA2<sup>27</sup> | 1550 ±225 | 45.00 ±4.20 | 2.90 |

**Figure 5.**
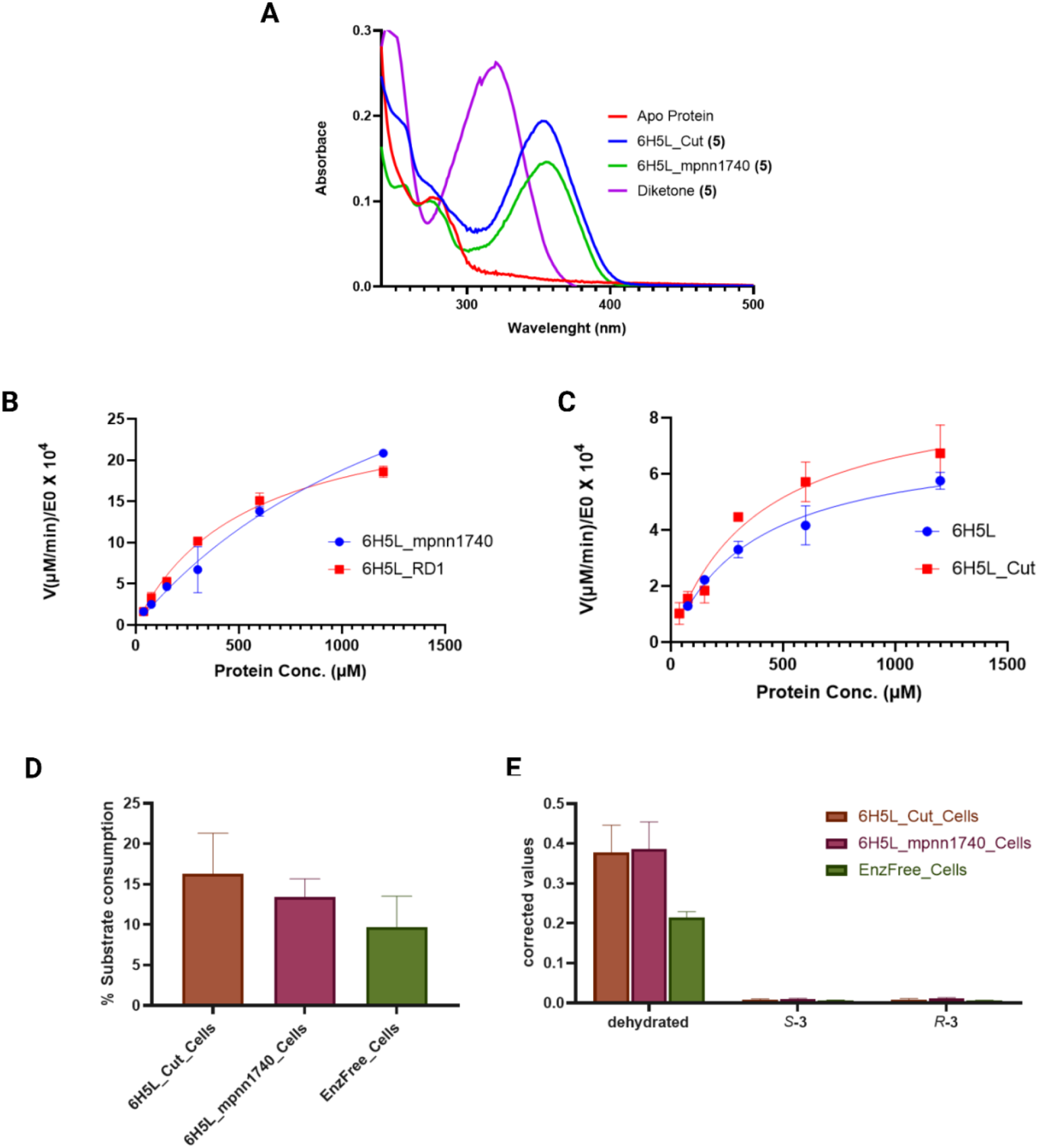
Activity assays of 6H5L_mpnn1740 and 6H5L_Cut. **A)** UV–Vis absorption spectra of diketone **5**, 6H5L apo-protein, and inhibitor-labeled proteins. Spectra show a new peak at 340 nm corresponding to vinylogous amide **6** following reaction with the diketone inhibitor **5. B), C)** Michaelis–Menten plots for retro-aldolase activity of all protein variants using methodol **3** as substrate. *E*_0_, enzyme concentration; *V*, initial reaction velocity. **D), E)** Substrate **1** consumption (%) and product formation (calculated from the area under the product peaks, *P*, relative to the internal standard, *IS*) from whole-cell aldol catalysis. Reactions were performed using 5 mM substrate **1** and 5 vol% substrate **2** in buffer (50 mM sodium phosphate, 150 mM NaCl, pH 7.5) containing 10 vol% acetonitrile (pH 7.5).

The truncated 6H5L_Cut design exhibited a *k*_cat_ / *K*_m_ of 2.38 M^−1^ min^−1^, marginally higher than that of the original 6H5L design (1.96 M^−1^ min^−1^) (Figure 5C, Table 2). While the *K*_m_ values were comparable, 6H5L_Cut showed a higher *k*_cat_ than the original 6H5L (Table 2). These results suggest that both termini of 6H5L_Cut remain open, forming two accessible binding pockets that retain comparable substrate binding while allowing a higher turnover rate (Figure 2A, Table 2).

#### Whole-cell biotransformation

Both redesigned variants, 6H5L_mpnn1740 and 6H5L_Cut, exhibited catalytic activity in whole-cell biotransformation of the aldol reaction between **1** (5 mM) and acetone **2** (in excess). Substrate consumption of **1** reached 16.3% and 13.4%, respectively, almost double that of enzyme-free cells (7.6%) (Figure 5D), in line with prior reports of 6H5L-mediated biotransformation^27^. Most of the aldol reaction product underwent dehydration, yielding an α,β-unsaturated ketone **4** at levels roughly double that of the enzyme-free controls, with only minor amounts of methodol **3** remaining (Figure 5E and S3).

## Discussion and Conclusions

The Rosetta-designed de novo asymmetric six-helical barrel (6H5L) provides a stable and catalytically active scaffold and is an ideal candidate scaffold for further computational design and engineering. However, it remains unclear how engineerable de novo beackbone-seuqneces pairs are. Arguably, a working de novo design represents already an optimal back bone-sequence pair and further modification could lead to detrimental effects in the structure or activity of such designed proteins. In this work, we used two different approaches to ask the fundamental question how engineerable this de novo scaffold is - ProteinMPNN-based sequence redesign and RosettaRemodel-based structural truncation.

Among the two ProteinMPNN-designed variants, both with low rosetta energies and high AF2 confidence (pLDDT > 90), only 6H5L_mpnn1740 (pI = 5.4) was successfully produced in soluble form, with a tenfold higher protein yield than the reference design 6H5L_RA1. In contrast, 6H5L_mpnn1409 (pI = 7.18) showed poor expression and multiple peaks in SEC. The two variants also differ in their sequence composition, charge distribution, and hydrophobicity, which may contribute to their different experimental behavior. However, with only two designs, it is difficult to draw a clear conclusion about the relationship between these properties and soluble expression. Remarkably though, both designs showed proper folding and thus highlight the high sequence diversity that can be achieved to stabilize the same de novo scaffold, a feature that is highly desirable for engineering purposes.

Despite the extensive sequence redesign, 6H5L_mpnn1740 retained the overall structure, high thermal stability, and retro-aldolase activity of the 6H5L scaffold. This shows that the scaffold can tolerate substantial sequence changes while maintaining its overall architecture and function. Compared with 6H5L_RA1, the variant showed a higher *k*_cat_ but also a higher *K*_m_, resulting in lower overall catalytic efficiency. The higher *K*_m_ may be related to changes in the unoptimized cavity 1 during the ProteinMPNN redesign, which could affect substrate binding.

Using RosettaRemodel, we also designed and experimentally validated the truncated construct 6H5L_Cut. The crystal structure (PDB 9R2B) closely matched the computational model, confirming that the six-helix barrel can be substantially shortened while preserving its overall architecture, the designed electrostatic interaction network formed by the charged residue ring in the barrel core, and the open hydrophobic channel. The truncated variant also retained high thermal stability and retro-aldolase activity. Compared with the original 6H5L, 6H5L_Cut showed comparable *K*_m_ values and a higher *k*_cat_, resulting in slightly higher catalytic efficiency. The increased flexibility observed at the helical termini may provide opportunities for further modifications of the barrel architecture for more efficient biocatalytic applications. In contrast, the second truncation design, 6H5L_Cut2, aggregated during expression and purification, suggesting that an average AF2 confidence score below 90 may not be sufficient to predict successful experimental behavior.

Both newly designed variants, 6H5L_mpnn1740 and 6H5L_Cut, retained high thermal stability and (retro)-aldolase activity in purified protein and whole-cell biotransformation experiments. The two design approaches modified the 6H5L scaffold in different ways, affecting protein production, structure, and catalytic properties. Together, these results show that DL-based and physics-based approaches can complement each other in the further engineering of de novo helical barrel biocatalysts, providing different routes to modify the scaffold while maintaining its stability and function, which could further expand their applications in biocatalysis and biotechnology.

## Methods

### Gene cloning

The designed protein variants, either ProteinMPNN or truncated versions, were synthesized as gene fragments for Gibson cloning^34^ into the pEThTEV vector, which incorporates an N-terminal His-Tag and a TEV cleavage site.

### Protein production

After confirming the cloned genes, they were expressed in *E. coli* BL21 Star (DE3) cells. A single colony was cultured overnight in LB medium supplemented with 50 mg/L kanamycin at 37 °C and 180 rpm. The pre-culture (10 mL) was inoculated into 1 L TB medium with 100 mg/L kanamycin and grown to an OD_600_ of ~ 0.6. Protein expression was induced with 1 mM IPTG, and the cells were incubated overnight at 18 °C and 180 rpm. Cells were harvested by centrifugation (4 °C, 4500 rpm, 30 min) and stored at -20°C. Cell lysis was performed by 10 min of sonication on ice in a buffer containing 20 mM sodium phosphate (pH 7.4), 500 mM NaCl, 10 mM imidazole, and 10 mM PMSF. The lysate was centrifuged (18,000 rpm, 45 min) to remove debris and insoluble protein.

### Protein purification

Protein variants were purified using Ni-NTA affinity and size-exclusion chromatography (SEC) with TEV-mediated His-tag cleavage. Lysates were loaded onto a Ni-NTA column, washed (20 mM sodium phosphate, pH 7.4, 500 mM NaCl, 30 mM imidazole), and eluted with the same buffer containing 250 mM imidazole. Concentrated proteins were rebuffered into Xtal buffer (20 mM Tris-HCl, 150 mM NaCl, pH 8) and incubated overnight with 15 mM TEV protease per 1 mg/mL protein. Cleaved proteins were collected from the Ni-NTA flowthrough and further purified by SEC (Superdex™ 200 increase for ProteinMPNN or Superdex™ 75 increase for truncated variants). Purification was monitored by SDS-PAGE, and protein concentrations were measured at 280 nm using a NanoDrop™ 2000.

### Circular dichroism spectroscopy (CD)

For secondary structure analysis, purified proteins were re-run on SEC using CD buffer (20 mM sodium phosphate, 150 mM sodium fluoride, pH 8.0). Eluted fractions were concentrated to 0.1 mg/mL and analyzed on a J-1500 CD spectrometer (JASCO, USA) using 0.5 mm quartz capillaries. Spectra were recorded between 190 and 260 nm, with temperature scans performed between 20 and 95 °C in 2 °C increments. CD signals were baseline-corrected using buffer blanks.

### Fluorescence ligand-binding experiment

Ligand-binding assays used 1,6-diphenylhexatriene (DPH) fluorescence dye, known for its sensitivity to hydrophobic environments, to validate correct barrel folding in *de novo* designed alpha-helix bundles^5^. Experiments were performed using a constant DPH concentration (1 µM) and varying protein concentrations from 0 to 100 µM in buffer (20 mM Tris, 150 mM NaCl, 1% DMSO, pH 8, at 25 °C). Measurements were conducted in 96-Well Black Flat Bottom Polystyrene Non-Binding Surface Microplates (Corning, Lowell, MA) using a BMG Labtech (Aylesbury, UK) Clariostar plate reader. After a 1-hour dark incubation with shaking at 300 rpm, fluorescence spectra were recorded with excitation at 350 ± 16 nm and emission between 380 and 602 nm. Dissociation constant (*K*_D_) values were determined by fitting the measured data to a one-site_total binding model using GraphPad Prism 8.

### Small-angle X-ray scattering (SAXS)

Both proteins were prepared for SAXS experiments by SEC in SAXS buffer (20 mM Tris-HCl, 150 mM NaCl, 2 mM TCEP, 3% glycerol, pH 8). Protein concentrations were adjusted to 3 mg/mL for 6H5L_mpnn1740 and 6 mg/mL for 6H5L_Cut. Measurements were conducted at the ESRF in Grenoble (France) on the BioSAXS beamline BM29. Scattering data were corrected by subtracting buffer scattering and then fitted against the theoretical scattering curves from AlphaFold2 models^16,35^ using the FoXS server to calculate fit quality (chi^2^ values)^32^.

### Crystallography

Crystallization drops were set up with commercial crystallization screens using the vapor diffusion method and employing a mosquito crystallization robot (SPT Labtech). Crystallization plates were incubated at 293 K. The protein concentration varied between 15-25 mg/mL in buffer containing 150mM NaCl, 1mM TCEP and 20mM Tris with pH 8. The drop volume was 270 nL, with a 1:1 protein and precipitant solution ratio. Crystallization drops were equilibrated against a reservoir containing 40 µL of precipitant solution. The crystals with the best diffraction were obtained from Morpheus A1. Obtained crystals were harvested from mother liquor with CryoLoops (Hampton Research) and briefly incubated with mother liquor containing 25% glycerol, followed by flash freezing in liquid nitrogen. Diffraction data was collected at 100 K at the ESRF in Grenoble (France) on beamline ID30-B. The collected data were processed using XDS^36^. Data resolution cutoffs were determined by pairef ^37^. Structure determination was performed by molecular replacement using PHASER^38^ with the design models as search templates. Models were automatically refined with PHENIX^39^ after manual model building using COOT^40^. Data collection and refinement statistics are summarized in Tables S2 and Figure 3.

### Inhibition reaction

The diketone inhibition assays (Figure 4B) were conducted in a buffer solution containing 50 mM disodium hydrogen phosphate, 150 mM NaCl, and 5% DMSO at pH 8. The reaction mixture included a protein concentration of 28 µM and 280 µM 1,3-diketone **5** (1- (6-methoxy-2-naphthalenyl)-1,3-butanedione; total volume = 1 mL. Following overnight incubation at 29 °C, reaction mixtures were reapplied to SEC to remove excess diketone inhibitor **5**. Protein fractions were collected, and UV-Vis spectra were recorded to compare the apo protein, inhibitor **5**, and product-bound proteins.

### Retro-aldolase activity

Kinetic assays were performed in 96-Well Black Flat Bottom Polystyrene Non-Binding Surface Microplates (Corning, Lowell, MA) using a plate reader (CLARIOstar, BMG Labtech). Initial reaction rates of the conversion of *rac*-4-hydroxy-4-(6-methoxy-2-naphthyl)-2-butanone **3** (methodol) to 6-methoxy-2-naphthaldehyde **1** (Figure 4A) were determined by measuring the fluorescence of **1** with an excitation wavelength of 330 nm and emission wavelength of 452 nm. All measurements were conducted at 29 °C in assay buffer (50 mM sodium phosphate, 150 mM NaCl, 7.5% DMSO, pH 8). The fluorescence signals were corrected by subtracting blank background reactions and converted to product concentrations using a calibration curve. Michaelis–Menten fitting within GraphPad Prism 8 was applied to determine catalytic rate constant (*k*_cat_) and Michaelis constant (*K*_m_) values.

### Whole-cell biotransformation

The activity assay and product characterization were performed as described previously^27^. *E. coli* BL21 Star (DE3) cells overexpressing protein variants were lyophilized, and their catalytic activity was tested. Cells containing empty plasmids served as controls. Lyophilized cells (15 mg) were resuspended in 300 µL reaction buffer (50 mM sodium phosphate, 150 mM NaCl, pH 7.5) for 20 min at 30 °C and 120 rpm, followed by the addition of 5 mM substrate 6-methoxy-2-naphthaldehyde **1** in 10 vol% acetonitrile and 5 vol% acetone (final volume 500 µL). Mixtures were incubated overnight at 30 °C and 120 rpm. Products were extracted with ethyl acetate (2 × 250 µL) containing 0.5 vol% acetophenone as internal standard (IS), dried over Na_2_SO_4_, and analyzed by HPLC (Chiralcel® OD-H, 250 × 4.6 mm, 5 µm; 5 µL injection). Results were corrected for buffer-catalyzed background reactions.

## Acknowledgements

We acknowledge the European Synchrotron Radiation Facility for provision of synchrotron radiation facilities, and we would like to thank the staff of the ESRF and EMBL Grenoble for assistance and support in using beamlines MASSIF-3, ID30B, and BM29. M.H thanks the University of Graz for funding. W.E. and G.O. were supported by the Austrian Science Fund (FWF) grant 10.55776/P30826 and by funding from the European Research Council through an ERC Starting Grant (HelixMold 802217). G.O. were supported by a FETOPEN project (ARTIBLED, 863170). This research was funded in whole, or in part, by the Austrian Science Fund (FWF) [10.55776/P30826 to GO].

## Author contributions

Conceptualization: W.E., G.O.; methodology: W.E, M.H, G.O.; Validation: W.E., B.G., D.S., M.C., M.H., A.B., G.O.; Formal analysis: W.E., M.H., G.O.; Investigation: W.E, B.G., D.S., M.C., M.A., M.H., G.O.; Resources: M.H., G.O.; Data Curation: W.E., G.O.; Writing: W.E., G.O.; Visualization: W.E., G.O; Supervision: M.H., G.O.; Project administration: G.O.; Funding acquisition: M.H., G.O.

## Data and materials availability

Supplementary Materials are available for this paper. Correspondence and requests for materials should be addressed to G.O.

## References

1. Wu, S., Snajdrova, R., Moore, J. C., Baldenius, K. & Bornscheuer, U. T. Biocatalysis: Enzymatic Synthesis for Industrial Applications. Angewandte Chemie International Edition 60, 88–119 (2021).

2. Jiang, L. et al. De Novo Computational Design of Retro-Aldol Enzymes. Science (1979) 319, 1387–1391 (2008).

3. Sterner, R., Merkl, R. & Raushel, F. M. Computational Design of Enzymes. Chem Biol 15, 421–423 (2008).

4. Zanghellini, A. et al. New algorithms and an in silico benchmark for computational enzyme design. Protein Science 15, 2785–2794 (2006).

5. Obexer, R. et al. Emergence of a catalytic tetrad during evolution of a highly active artificial aldolase. Nat Chem 9, 50–56 (2017).

6. Burton, A. J., Thomson, A. R., Dawson, W. M., Brady, R. L. & Woolfson, D. N. Installing hydrolytic activity into a completely de novo protein framework. Nat Chem 8, 837–844 (2016).

7. Garrabou, X., Macdonald, D. S. & Hilvert, D. Chemoselective Henry Condensations Catalyzed by Artificial Carboligases. Chemistry – A European Journal 23, 6001–6003 (2017).

8. Das, R. & Baker, D. Macromolecular Modeling with Rosetta. Annu Rev Biochem 77, 363–382 (2008).

9. Fleishman, S. J. et al. RosettaScripts: A Scripting Language Interface to the Rosetta Macromolecular Modeling Suite. PLoS One 6, e20161 (2011).

10. Ferruz, N., Schmidt, S. & Höcker, B. ProtGPT2 is a deep unsupervised language model for protein design. Nat Commun 13, 4348 (2022).

11. Wicky, B. I. M. et al. Hallucinating symmetric protein assemblies. Science (1979) 378, 56–61 (2022).

12. Anishchenko, I. et al. De novo protein design by deep network hallucination. Nature 600, 547–552 (2021).

13. Wang, J. et al. Scaffolding protein functional sites using deep learning. Science (1979) 377, 387–394 (2022).

14. Singer, J. M. et al. Large-scale design and refinement of stable proteins using sequence-only models. PLoS One 17, e0265020 (2022).

15. Sumida, K. H. et al. Improving Protein Expression, Stability, and Function with ProteinMPNN. J Am Chem Soc http://www.10.1021/jacs.3c10941 (2024) doi:10.1021/jacs.3c10941.

16. Guo, H.-B. et al. AlphaFold2 models indicate that protein sequence determines both structure and dynamics. Sci Rep 12, 10696 (2022).

17. Baek, M. et al. Efficient and accurate prediction of protein structure using RoseTTAFold2. biorxiv 2023.05.24.542179 10.1101/2023.05.24.542179 (2023).

18. Lin, Z. et al. Evolutionary-scale prediction of atomic-level protein structure with a language model. Science (1979) 379, 1123–1130 (2023).

19. Wu, R. et al. High-resolution de novo structure prediction from primary sequence. biorxiv 2022.07.21.500999 10.1101/2022.07.21.500999 (2022).

20. Trost, B. M. & Brindle, C. S. The direct catalytic asymmetric aldol reaction. Chem Soc Rev 39, 1600 (2010).

21. Yamashita, Y., Yasukawa, T., Yoo, W.-J., Kitanosono, T. & Kobayashi, S. Catalytic enantioselective aldol reactions. Chem Soc Rev 47, 4388–4480 (2018).

22. Althoff, E. A. et al. Robust design and optimization of retroaldol enzymes. Protein Science 21, 717–726 (2012).

23. Albery, W. J. & Knowles, J. R. Evolution of enzyme function and the development of catalytic efficiency. Biochemistry 15, 5631–5640 (1976).

24. Kipnis, Y. et al. Design and optimization of enzymatic activity in a de novo β-barrel scaffold. Protein Science 31, (2022).

25. Ożga, K. & Berlicki, Ł. Miniprotein-Based Artificial Retroaldolase. ACS Catal 12, 15424–15430 (2022).

26. Braun, M. et al. Computational design of highly active *de novo* enzymes. bioRxiv 2024.08.02.606416 (2024) doi:10.1101/2024.08.02.606416.

27. Elaily, W. et al. Computational design of a thermostable de novo biocatalyst for whole cell biotransformations. bioRxiv 2024.10.07.617055 (2024) doi:10.1101/2024.10.07.617055.

28. Huang, P.-S. et al. High thermodynamic stability of parametrically designed helical bundles. Science (1979) 346, 481–485 (2014).

29. Albanese, K. I. et al. Rationally seeded computational protein design of ?-helical barrels. Nat Chem Biol 20, 991–999 (2024).

30. Huang, P.-S. et al. RosettaRemodel: A Generalized Framework for Flexible Backbone Protein Design. PLoS One 6, e24109 (2011).

31. Thomas, F. et al. De Novo -Designed α-Helical Barrels as Receptors for Small Molecules. ACS Synth Biol 7, 1808–1816 (2018).

32. Schneidman-Duhovny, D., Hammel, M., Tainer, J. A. & Sali, A. Accurate SAXS Profile Computation and its Assessment by Contrast Variation Experiments. Biophys J 105, 962–974 (2013).

33. Tanaka, F., Fuller, R., Shim, H., Lerner, R. A. & Barbas, C. F. Evolution of Aldolase Antibodies in Vitro : Correlation of Catalytic Activity and Reaction-based Selection. J Mol Biol 335, 1007–1018 (2004).

34. Richter, F., Leaver-Fay, A., Khare, S. D., Bjelic, S. & Baker, D. De Novo Enzyme Design Using Rosetta3. PLoS One 6, e19230 (2011).

35. Gibson, D. G. et al. Enzymatic assembly of DNA molecules up to several hundred kilobases. Nat Methods 6, 343–345 (2009).

36. Mirdita, M. et al. ColabFold: making protein folding accessible to all. Nat Methods 19, 679–682 (2022).

