## Supplementary Materials for "Probing the sequence variability tolerance in a de novo α-helical barrel biocatalyst"

### For

**Content:**

- Supplementary Figures S1 – S3
- Supplementary Table S1 & S2

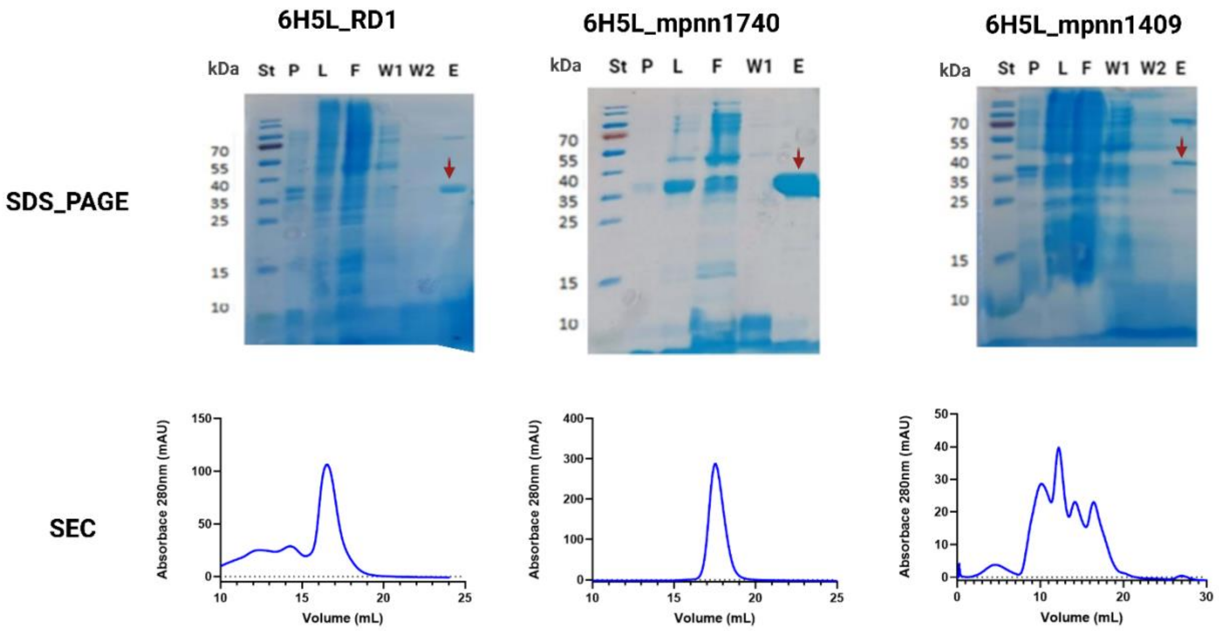


**Figure S1.** SDS-PAGE analysis of protein samples following affinity purification. St, protein ladder (Thermo Scientific Prestained Protein Ladder 10–180 kDa); P, cell pellet; L, cell lysate; F, flow-through; W1 and W2, wash fractions 1 and 2, respectively; E, elution fraction. Size-exclusion chromatography (SEC) profiles of the ProteinMPNN designs 6H5L_mpnn1740 and 6H5L_mpnn1409, as well as the input design 6H5L_RA1.

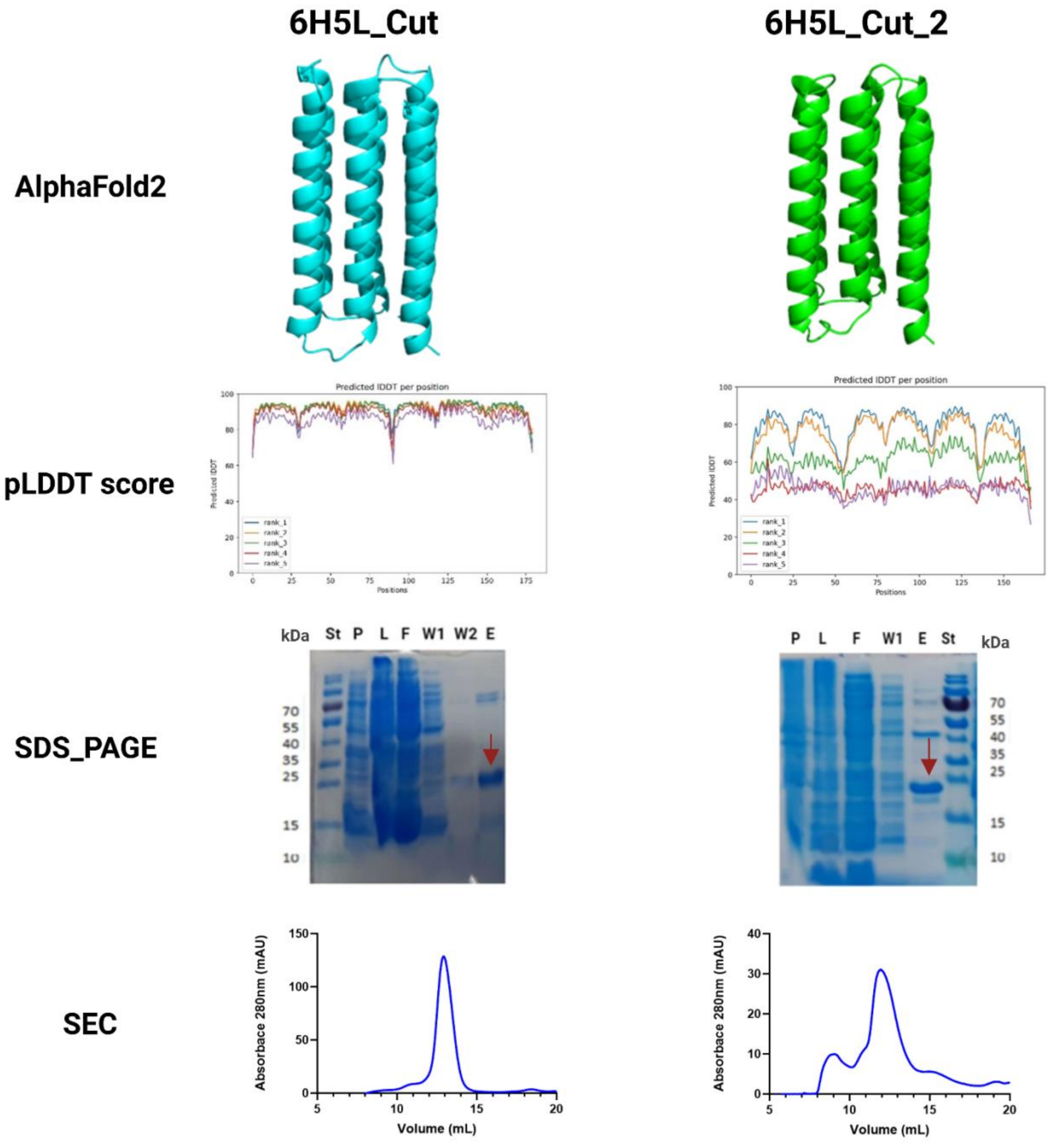


**Figure S2.** AlphaFold2-predicted structures with per-residue pLDDT scores, SDS-PAGE analysis of protein samples following affinity purification, and size-exclusion chromatography (SEC) profiles of the truncated designs 6H5L_Cut and 6H5L_Cut2. St, protein ladder (Thermo Scientific Prestained Protein Ladder 10–180 kDa) ; P, cell pellet; L, cell lysate; F, flow-through; W1 and W2, wash fractions 1 and 2, respectively; E, elution fraction.

**Table S1**. Amino acid sequences and theoretical pIs of all 6H5L protein variants.

| Protein | Sequence | Theoretical pI |
|---|---|---|
| **6H5L** | TCEVVKKILYMAKKLVEKKKEVVKKILKMAKELVEKKKEVVYKAL EAAEKGLDTKKIAKLLLEMLEHELELAEKIAKLLLEMLEEELKLAE KIAKLMLESGISEEVVKKILYMAKKLVEKKKEVVKKILYMAKELVE KKKEVVKKALEAAEKGLDTKKIAKLLLEMLEHELELAEKIAKLLLE MLEEELKLAEKIAKLMLESGISEEVVKKILKMAKKLVEKKKEVVKK ILYMAKELVEKKKEVVKKALEAAEYGLDTKKIAKLLLEMLEHELEL AEKIAKLLLEMLEEELKLAEKIAKLMEEQCK | 9.05 |
| **6H5L_mpnn 1740** | LEEVVKKQLELAKELVEKQKKVVEEELKLAKELAELKREVVVKA QELAQAGKSTEPIAELALKYLDKLLEAAKKEAELKLKALEEQLKA AVEEYKLFKEAGVESKVVEKILELAKKVVEEQKKVVEKELELAKK LIEKKKKVAKEALEYAQKGKSTEEIAKGLLEMAEEERKAAKEAAE LKLKALEKQEKAAEKIYELFKEEGVESEVVEEALKLAKEAVELQK KVVEEELKLAEKYIEKVKEVVKKAQEVAQAGKSTQIIADLLLEYEK FQVEIAEKEAKLKLEALEKQLEFAKKIAELFKKEAE | 5.49 |
| **6H5L_mpnn 1409** | LGAAVEAQLKLSEELVAKQRDVVKKQLELDKALAKEKLKVVEEA HKLAKEGKGTEEITKLALEYLKKLLEAEKKEAELWLSALAAQKAA AEEKYKLFLAAGVKSKAVPAQLALSKAEVAAQEDAVQKQLELDK EVVKKKEEVAKEAHKAAKKGKGTEPIAEGLLEMAKFEEKAEEEQ ANLWLSVLDAQEASAVQKALLFLAAGVKSEIVPKELELSKQLVAA QKAVVKKQLELDKKYVEKVKEVVKYAHEVAKEGKGTEEIAKKLK EYVEFQVKIEEQEAAAWLAALAKQAAAAKEKAELFKKAAE | 7.18 |
| **6H5L_Cut** | TCEVVKKILYMAKKLVEKKKEVVKKIEKGQEDPATLLLEMLEEEL KLAEKIAKLMLESGISEEVVKKILYMAKKLVEKKKEVVKKIEKGGT DPAELLLEMLEEELKLAEKIAKLMLESGISEEVVKKILKMAKKLVE KKKEVVKKIAKDRKNAAKLLLEMLEEELKLAEKIAKLMEEQCKGS GW | 9.09 |
| **6H5L_Cut2** | PEEVVKKILYMAKKLVEKKKKASKTSKSAALETLEEELKLAEKIAK LYKAGGLDGLPTQKILYMAKKLVEKKKEVVIYGGGDSALLEMLE EELKLAEKIAAEWARKGLSDSVAEEILKMAKKLVEKKKEVVKKSE TGVDDEMLEEELKLAEKIAKLMEEQCK | 8.72 |

**Table S2.** Crystallographic data collection and refinement statistics for 6H5L_Cut1.

| **Parameter** | **Value** |
|---|---|
| Protein | 6H5L_Cut1 |
| Wavelength (Å) | 0.8731 |
| Resolution range (Å) | 37.28–2.1 (2.15–2.1) |
| Space group | C 1 2 1 |
| Unit-cell parameters (Å, °) | 149.4 33.27 102.71 90 133.13 90 |
| Total reflections | 112936 (5227) |
| Unique reflections | 61200 (3322) |
| Multiplicity | 1.8 (1.6) |
| Completeness (%) | 99.59 (99.87) |
| Mean I/σ(I) | 3.62 (-0.12) |
| Wilson B-factor | 30.81 |
| R-merge | 0.1157 (-10.9) |
| R-meas | 0.1614 (-15) |
| R-pim | 0.112 (-10.23) |
| CC1/2 | 0.996 (-0.0237) |
| CC* | 0.999 (-0.22) |
| Reflections used in refinement | 21959 (1540) |
| Reflections used for R-free | 1600 (119) |
| R-work | 0.2665 (0.2761) |
| R-free | 0.3074 (0.3176) |
| Number of non-hydrogen atoms | 2781 |
| Macromolecules | 2551 |
| Ligands | 39 |
| Solvent | 191 |
| Protein residues | 339 |
| RMS(bonds) | 0.002 |
| RMS(angles) | 0.40 |
| Ramachandran favored (%) | 98.78 |
| Ramachandran allowed (%) | 1.22 |
| Ramachandran outliers (%) | 0.00 |
| Rotamer outliers (%) | 1.15 |
| Clashscore | 2.77 |
| Average B-factor | 37.55 |
| Macromolecules | 37.01 |
| Ligands | 48.38 |
| Solvent | 42.53 |

HPLC chromatograms of the aldol reaction catalyzed by whole-cell biocatalyst.

**Blank**

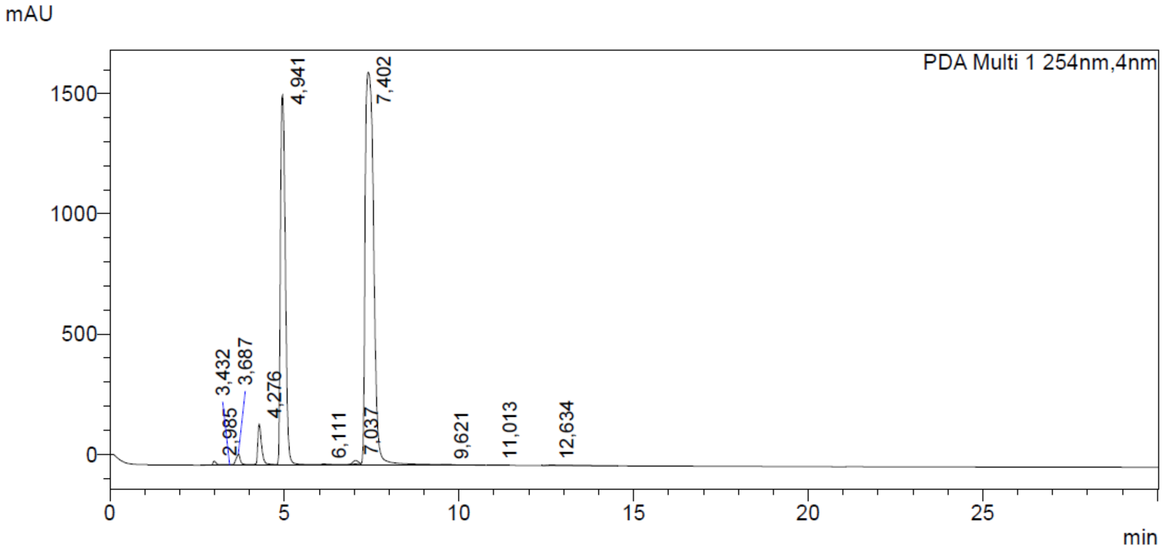


**Enzyme free cells**

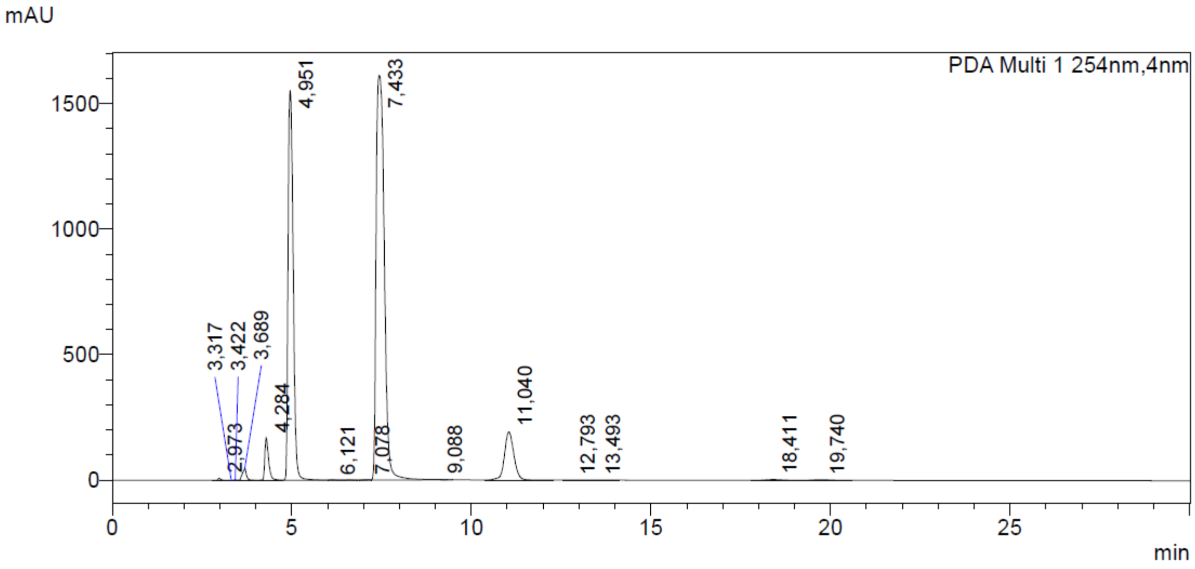

**6H5L_Cut cells**

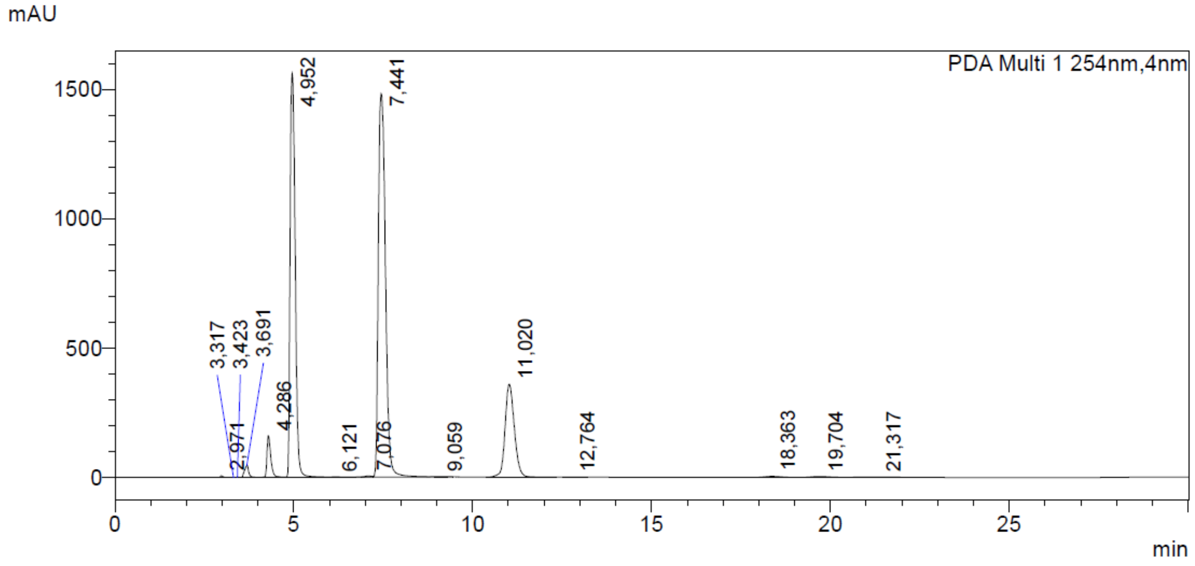


**6H5L_mpnn1740 cells**

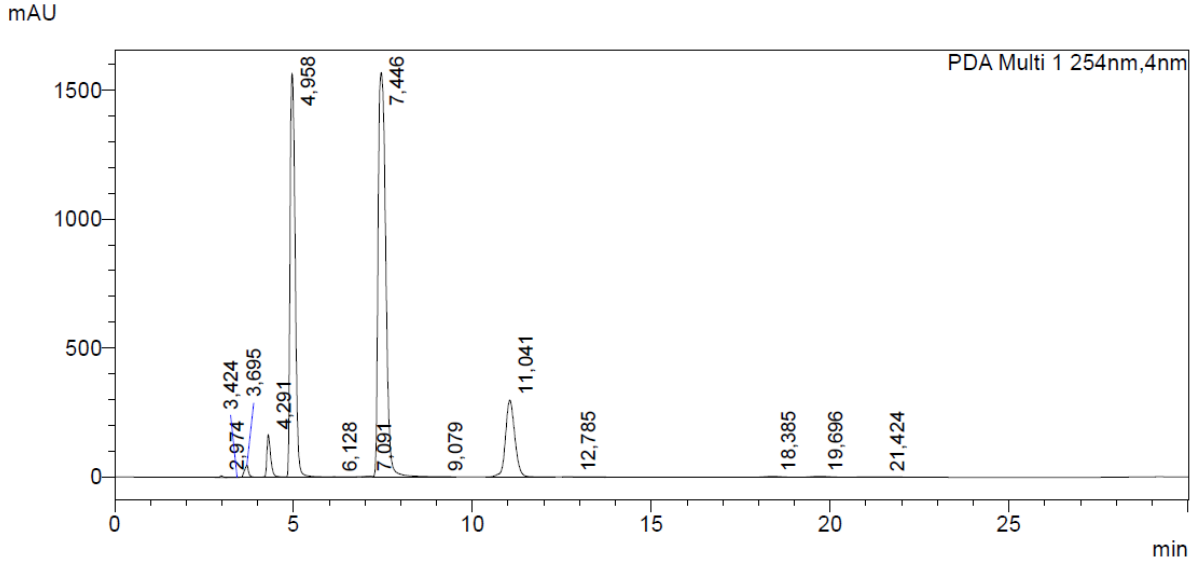


**Figure S3.** HPLC chromatograms of aldol reactions, including blank samples and reactions catalyzed by enzyme-free cells, 6H5L_Cut cells, and 6H5L_mpnn1740 cells (5 mM aldehyde **1** and 5 vol% acetone in buffer: 50 mM sodium phosphate, 150 mM NaCl, pH 7.5, 10% MeCN). The chromatograms show the internal standard (acetophenone) at 4.9 min, substrate **1** at 7.4 min, dehydration product **4** at 11.0 min, and methodol enantiomers (*R*)-**3** at 18.3 min and (*S*)-**3** at 19.6 min.
